# Preparing to act follows Bayesian inference rules

**DOI:** 10.1101/2024.08.16.608232

**Authors:** Luca Tarasi, Chiara Tabarelli de Fatis, Margherita Covelli, Giuseppe Ippolito, Alessio Avenanti, Vincenzo Romei

**Affiliations:** Department of Psychology, University of Bologna and Centre for Studies and Research in Cognitive Neuroscience, University of Bologna, Cesena, Italy; Laboratory of Cognitive Neuroscience, University of Udine, Udine, Italy; Universidad Antonio de Nebrija, Madrid, Spain

**Author notes:** Corresponding Author: Vincenzo Romei, Centro Studi e Ricerche in Neuroscienze Cognitive, Dipartimento di Psicologia, Alma Mater Studiorum – Università di Bologna, Campus di Cesena, via Rasi e Spinelli, 176. 47521 Cesena, Italy.

## Abstract

Predictive brain theories suggest that human brain sets-up predictive models to anticipate incoming sensory evidence. Recent studies demonstrated these models to be integrated already in sensory areas, shaping even perceptual outcomes. Here, we hypothesized that this integration process informs the entire functional hierarchy, thus scaling all the way down to the motor system. Operationally, we propose that cue-oriented cortico-spinal excitability (CSE) modulation serves the pre-activation of motor representations aligned to prior-congruent decisions. To this end, 62 participants completed a probabilistic discrimination task while we delivered bilateral single-pulse TMS over the two primary motor cortices (M1s) and recorded motor-evoked potentials (MEPs) to assess motor excitability associated to prior-congruent vs. incongruent actions separately encoded by the two hands. Our findings revealed that prior expectations shaped CSE well before action execution, predominantly by inhibiting the M1 cortex coding for the prior-incongruent action. Importantly, this physiological modulation underpinned the prior-induced bias in participants’ choices, highlighting the link between motor preparatory modulation and actual decision-making. Furthermore, we observed significant interindividual variability in prior-driven CSE modulations, revealing two distinct predictive strategies: the *believers’* style, who heavily rely on prior, and the *empiricists* one, who downplay its role, maintaining CSE level mostly unbiased. Crucially, autistic and schizotypal traits drove these differences in prior-driven motor strategy, with *believers* characterized by schizotypal traits, whereas *empiricists* displaying autistic-like features. These results demonstrate how predictive models are integrated into action representations and highlight the role of prior-driven CSE modulations as a potential marker able to intercept interindividual differences in predictive styles.

## Introduction

Prior expectations shape perceptual outcomes, modulating response rates (Bang and Rahnev, 2017), reaction times (Mulder et al., 2012), confidence (Tarasi et al., 2024) and metacognition (Sherman et al., 2015). Here, we investigated the physiological mechanisms by which the motor system encodes the priors influencing perceptual decision-making. Recent studies have demonstrated that prior knowledge about upcoming stimuli affects perceptual areas activity, with this modulation correlating with prior-dependent behavioral change (Kloosterman et al., 2019; Samaha et al., 2015; Tarasi et al., 2022a). We propose that a crucial mechanism through which prior information could influence behavior is by pre-activating the motor representation aligned with the prior-congruent decision. Motor areas activity is crucial in action preparation, with this activity modulated by the probability of impending actions (Bestmann et al., 2008) and decision-related variables such as reward anticipation (Bundt et al., 2016; Klein et al., 2012). However, while most studies have investigated the motor preparation of actions that have already been selected, the role of prior expectations about stimulus identity in shaping preparatory motor cortex excitability has not been thoroughly evaluated. Furthermore, the specific mechanism by which priors could be integrated within the motor system remains unknown. One hypothesis is that motor cortex region coding for the expectation-congruent response increases its excitability (*prior-excitation hypothesis*) to facilitate prior integration. Alternatively, this process might depend on the inhibition of the motor cortex region encoding the prior-incongruent response (*prior-inhibition hypothesis*).

To test our hypotheses, 62 participants performed a Random-Dot Motion Task task in which shifts in decision bias were experimentally induced within participants using a probabilistic cue that signalled the likelihood of left- or rightward movement. Analysis through Signal Detection Theory (SDT) and the Drift Diffusion Model (DDM) revealed that these expectations biased participants’ decisions towards the anticipated direction, such as increasing leftward reports when the cue suggested left movement. To substantiate a neurophysiological mechanism for this effect, we demonstrated that prestimulus motor-evoked potentials (MEPs) elicited with transcranial magnetic stimulation (TMS) on the two M1 areas where higher when the action was congruent with prior expectations and lower MEPs when incongruent, suggesting participants prepare their prestimulus motor responses based on prior information. This suggests that the brain’s preparatory processes are influenced by expectations before the actual stimulus is encountered, highlighting the profound impact of prior information on action planning. Additionally, individual differences in MEPs modulations aid us in revealing two predictive strategies: the believers one, who heavily rely on prior cues and was associated to schizotypal personality, and the empiricist’ strategy, who show less reliance and were characterized by higher autistic-like traits.

All in all, these findings identify a neurophysiological mechanism through which intentional modulation of motor excitability is applied to strategically shape perceptual decisions in probabilistic scenarios and reveal how the implementation of this mechanism exhibits significant variability depending on individual differences in the predictive strategies employed.

## Results

Human participants (N = 62) performed a Random Dot Motion (RDM) discrimination task, where they indicated the perceived direction of movement (i.e., rightward or leftward) (Fig.1A). To manipulate expectations about the upcoming stimulus, a probabilistic cue was presented at the centre of the screen before the stimulus appeared. Following a delay, the random-dot stimulus was shown at the centre of the screen for 400 ms. Presenting the stimulus centrally controlled for attentional modulation, ensuring that participants focused on the centre of the screen to perform the task. Responses required an index finger abduction to press a key on a reversed keyboard: F12 with the right hand for signalling rightward movement and F5 with the left hand for signalling leftward movement. Throughout the task, electromyographic (EMG) activity from the first dorsal interosseous (FDI) muscle in both hands (which acts as the agonist in index finger abduction) was recorded, and single-pulse TMS was administered to both the right and left M1 at four different timings (−700 ms, −400 ms, −100 ms, +200 ms relative to stimulus onset) at 110% of the previously determined resting motor threshold (rMT).

**Figure 1.**
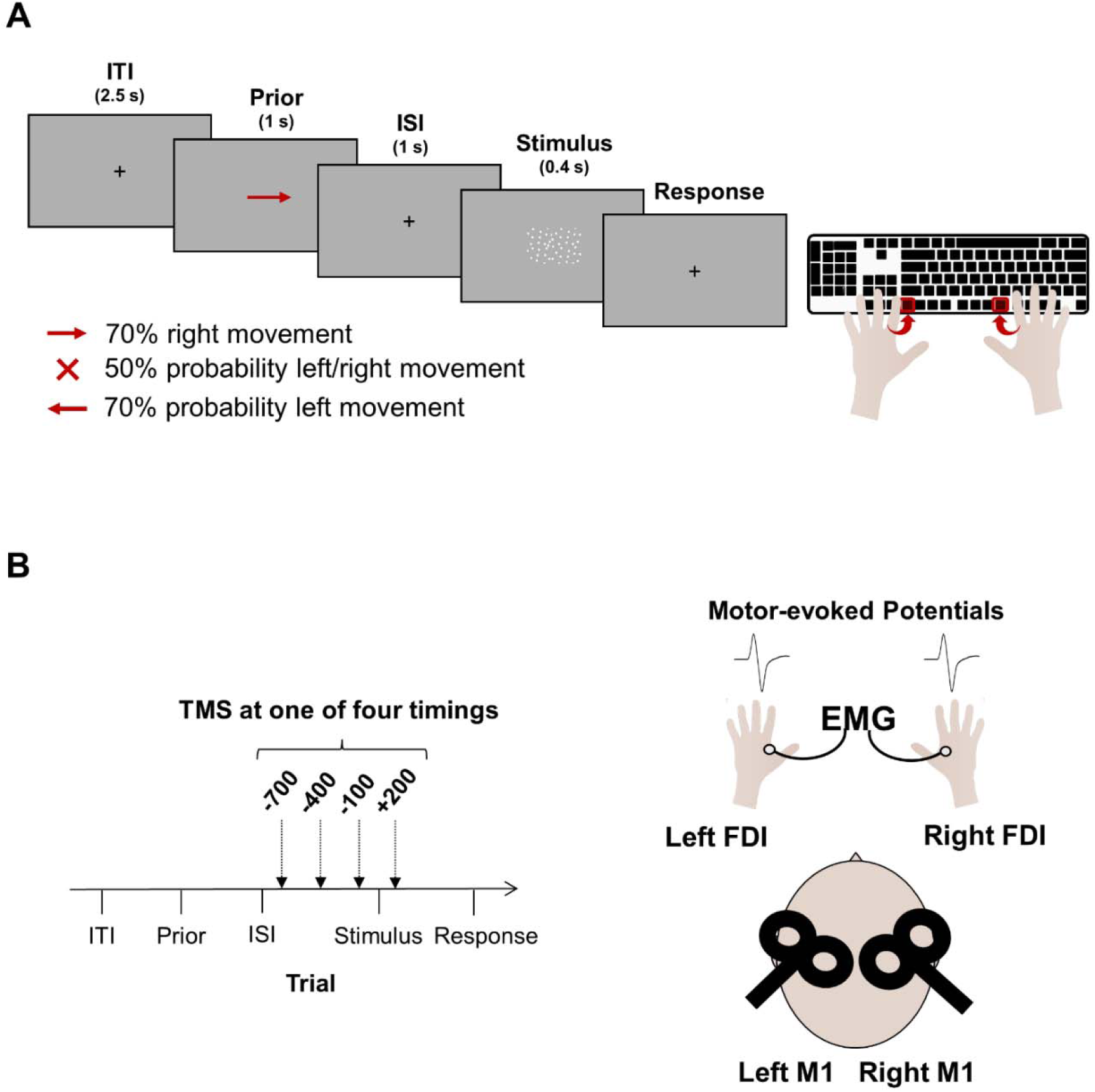
Experimental design. **A.** Behavioral and physiological data were collected during a motion discrimination task. Each trial started with a fixation cross, followed by a probabilistic cue indicating the probability of rightward vs. leftward movement. After a delay of 1 second, the dot motion stimulus appeared at the center of the screen for 0.4s. Cues were either informative (leftward vs. rightward arrow indicating 70% probability of leftward vs. rightward movement) or uninformative (a cross indicating 50% probability of either direction). Participants had to indicate the perceived direction of movement by making an abduction movement with the corresponding hand (i.e., right for rightward movement and left for leftward movement). **B.** In each trial, bilateral single-pulse TMS was delivered on right and left M1 at one of four timings (−700, −400, −100, +200 ms relative to stimulus onset). Motor-evoked potentials were recorded from right and left FDI.

Signal Detection Theory (SDT) analysis (Fig.2A) revealed a significant cue effect on response criterion (i.e., the tendency to respond “right” or “left”; F_2,122_=37.96, p<0.001) but not on sensitivity (F_2,122_=1.18, p=0.31). Specifically, in trials preceded by the left vs. right cue, participants showed a bias in indicating a leftward vs. rightward responses. Thus, our findings corroborate previous studies showing that expectations are integrated in the decision-making process, making the prior-congruent option more likely to be chosen (Bang and Rahnev, 2017; Tarasi et al., 2022a; Wyart et al., 2012).

**Figure 2.**
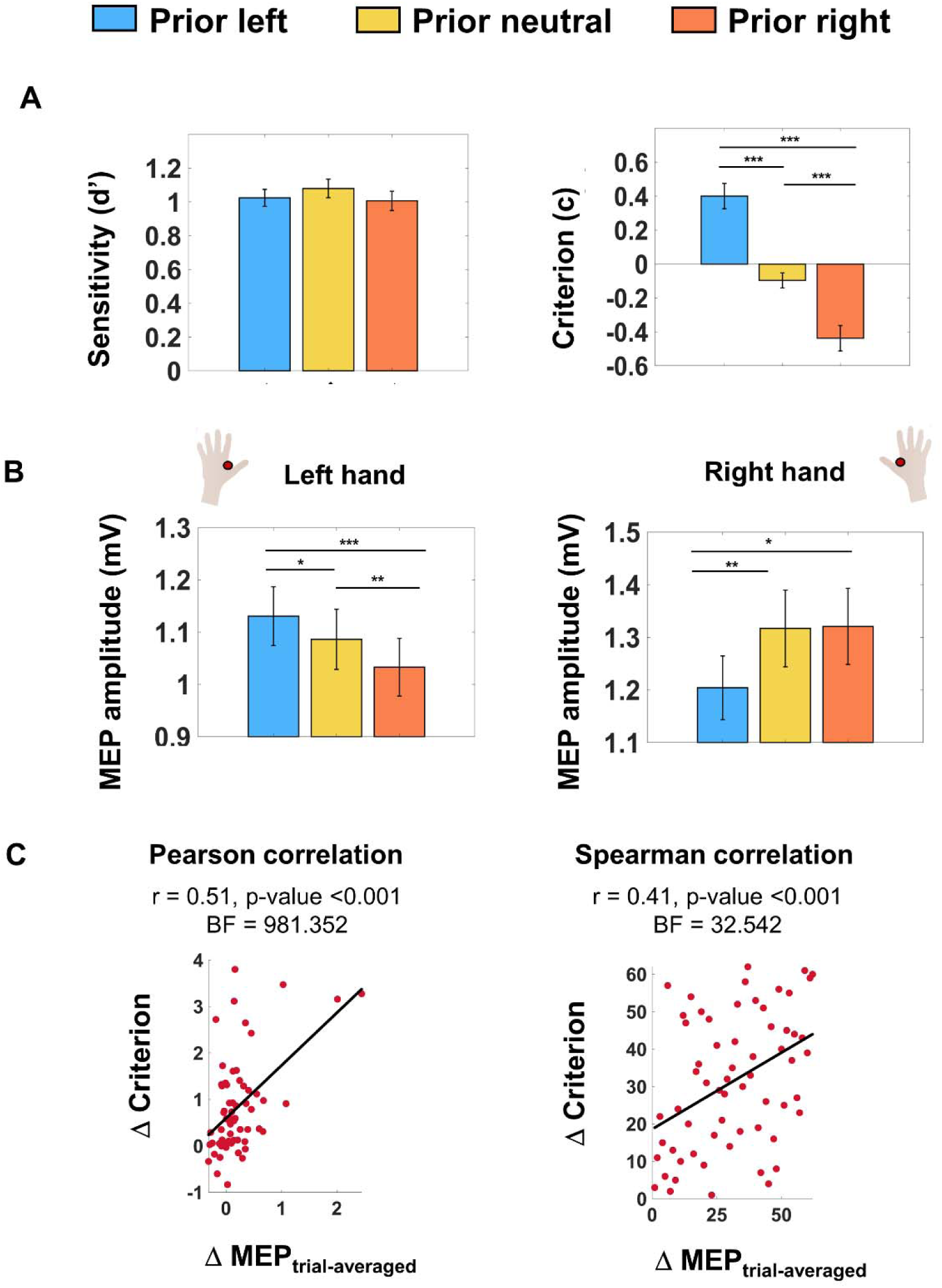
Prior-induced MEPs modulation correlates with decisional bias. **A.** SDT indices for Sensitivity (d’) and Criterion (c) are represented separately for trials preceded by leftward, rightward and neutral cues. Probabilistic cues did not influence sensitivity; however, they did affect criterion. Positive vs. negative criterion values indicate a tendency to respond left vs. right, therefore indicating a bias in responding left with the leftward cue (c_leftward_ = 0.40 ± 0.07) and a bias in responding right with the rightward cue (c_rightward_ = −0.44 ± 0.08). All comparisons revealed significant differences between the criterion adopted in the three conditions (c_leftward_ vs c_rightward_: t=6.33; c_leftward_ vs c_neutral_: t= 6.06; c_rightward_ vs c_neutral_: t=5.53; all p<0.001). Data are represented as mean ± SEM; *p<0.05, **p<0.01, ***p<0.001. **B.** Mean MEPs amplitude (mV) recorded from left and right hand are represented separately for trials preceded by the three probability cues. Prior information modulated MEP activity accordingly: the left hand showed higher vs. lower activity after the leftward vs. rightward cue, while the right hand showed the opposite pattern, thus indicating a congruency effect between hand and prior. With the neutral cue, activity in the left hand was in between the congruent and incongruent conditions, while activity in the right hand was the same as with the rightward cue. Error bars represent SEM; *p<0.05, **p<0.01, ***p<0.001. **C.** Correlation analysis between Δ Criterion and Δ MEP_trial-averaged_ showed a significant positive relationship between prior-dependent modulations of response criterion and MEPs amplitude. Thus, a greater prior-induced shift in MEPs amplitude is mirrored by more substantial changes in decision behavior.

Then, we investigated whether prior information could shape Cortico-Spinal Excitability. To this end, EMG activity was recorded from the first dorsal interosseous muscle of both hands during task execution. In each trial, the two primary motor cortices (i.e., left and right M1) were stimulated using single-pulse TMS. Pulses were delivered at different timing relative to the onset of the stimulus (i.e., −700, −400, −100, +200 ms; Fig.1B) in randomized order across trials. This protocol allowed to assess potential prior-related changes in MEPs amplitude. Indeed, in each prior informative trial, one motor cortex encoded the prior-congruent response (e.g., right M1 in prior left trials), while the other encoded the prior-incongruent response (e.g., left M1 in prior left trials).

The analysis showed a significant interaction between hand and prior (F_2,122_=12.41, p<0.001), indicating that the prior-dependent modulation of MEPs differed between the two hands (Fig.2B). Post-hoc analyses revealed a congruency effect between hand and cue: in the right hand, MEPs were higher (1.32±0.07) vs. lower when a rightward vs. leftward cue (1.20±0.06; t_61_=-2.68, p=0.01;BF=3.59) was presented. After the neutral cue, MEPs were higher (1.32±0.07) compared to the leftward condition (t_61_=-3.18; p=0.002; BF=12.44), while no difference emerged when considering the rightward trials (t_61_=-0.20; p=0.844; BF=0.14). As for the left hand, MEPs were higher (1.13±0.06) when a leftward cue was presented relative to when a rightward (1.03±0.6, t_61_=4.20, p<0.001;BF=239.49) or neutral cue (1.09±0.06; t_61_=2.54; p=0.014; BF=2.64) were presented. Additionally, higher MEPs were observed in neutral trials compared to the right cue condition (t_61_=3.18, p=0.002;BF=12.64). Overall, regardless of the hand considered, motor excitability was lower for prior-incongruent compared to congruent and neutral actions. We also observed a diametric effect, heightening the excitability in the prior-congruent vs. neutral condition, with this effect being specific for the left hand, suggesting the presence of complementary mechanism for integrating prior in the non-dominant motor pathway. Importantly, these findings hold valid even when removing the condition in which pulses were delivered after stimulus presentation (See SI). Furthermore, the prior-driven MEP modulations were unrelated to TMS timing, as it did not interact with other factors. Finally, we proved a strong association between the effect of prior information at the behavioral and physiological level. Specifically, the more participants shifted their motor cortex excitability in a prior-dependent fashion, the more they relied on prior information in their decision-making strategy (Fig.2C; Pearson = 0.51, p<0.001, BF=981.35, Pearson_skipped_=0.31, CI=[0.07;0.50]; Spearman=0.41, p=0.001, BF=32.54, Spearman_skipped_=0.34, CI=[0.06;0.55]).

To provide conclusive and robust evidence on the role played by the motor system in integrating predictive information, we employed a Drift Diffusion Model (DDM) approach to investigated whether trial-by-trial modulation of MEPs amplitude was able to modulate the prior-driven bias induction. We fit two different DDM models (See methods and SI), both converging in suggesting a strong relationship between trial-by-trial variation in MEPs and starting point shift (i.e., how biased participants are) (Fig. 3). The first model indicated that trials in which MEPs of the prior-congruent hand (e.g., right hand in right prior condition) were higher than MEPs recorded in the prior-incongruent hand (e.g., left hand in right prior condition) were associated with a stronger bias in responding in accordance with the expectation-like information provided by the cue.

**Figure 3.**
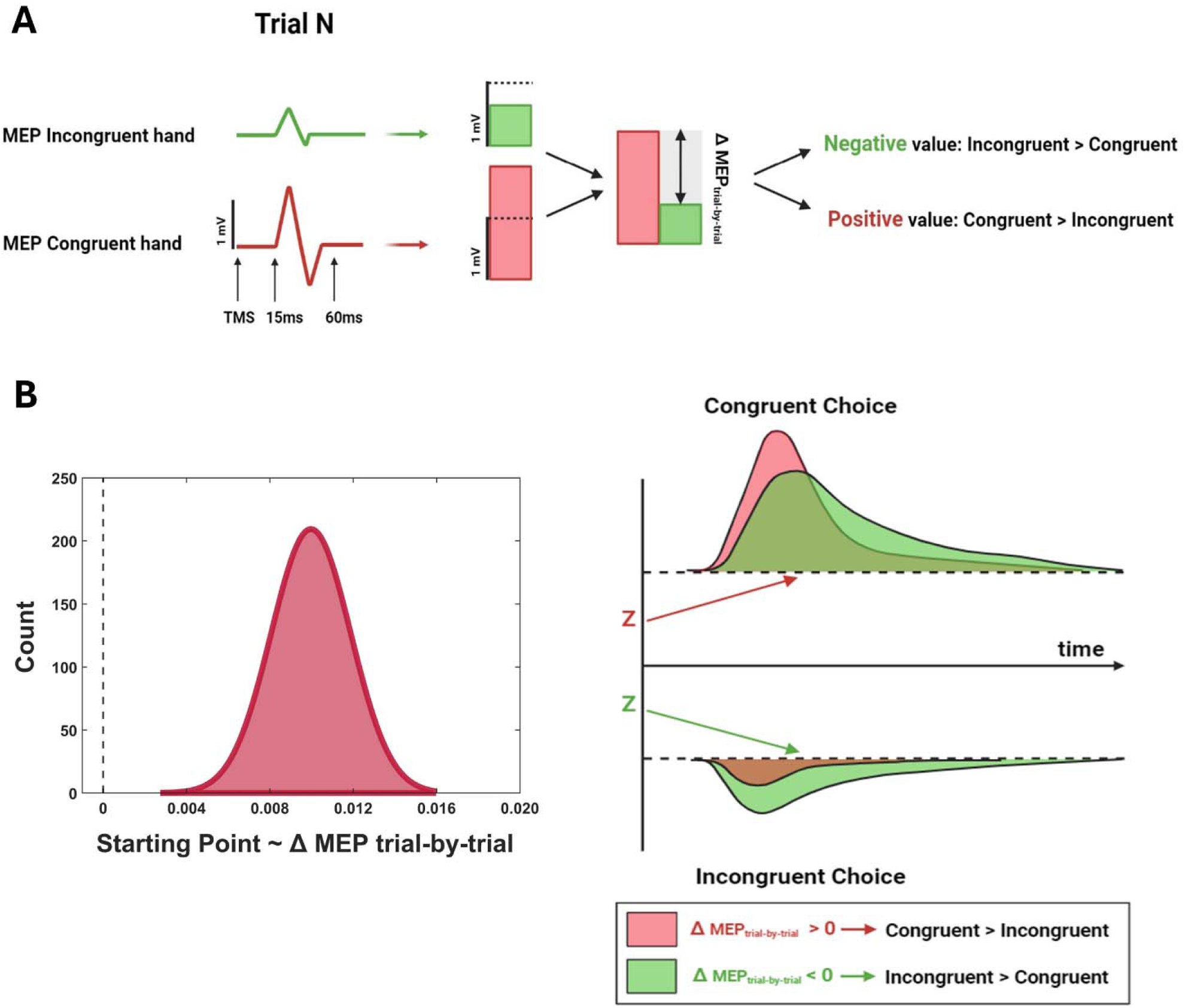
Trial-by-trial analysis. **A.** We setup a DDM to determine whether the trial-by-trial variability between congruent and incongruent MEPs influenced participants towards congruent or incongruent decisions. We coded z-scored MEPs based on its congruency. MEPs were considered congruent if they originated from the hand corresponding to the prior-congruent response, and vice versa for incongruent MEPs. Then, we calculated the difference between congruent and incongruent MEP (ΔMEP_trial-by-trial_) to derive a single index suitable for fitting the DDM. **B.** DDM indicated the significant effect of the MEP_trial-by-trial_ parameter. The coefficient is positive, means that a positive vs. negative ΔMEP_trial-by-trial_ value was associated to a higher vs. lower bias in responding in accordance with the expectation-like information provided by the cue.

The second model indicated that the highlighted effect holds valid even when removing the congruency effect, showing that trials in which the MEPs of the right (vs. left) hand were relatively larger in amplitude than those of the left (vs. right) hand, were associated with a stronger rightward (vs. leftward) decisional bias (Fig.S2). Crucially, this effect interacted with the provided priors. Specifically, while this significant effect persisted under conditions where the prior was informative (i.e., right and left), MEPs fluctuations did not predict the shift in the starting point parameter in the condition where the prior was uninformative. This suggests that the trial-by-trial effect is not merely a stochastic mechanism, where the decision is shifted based on which M1 area exhibits a stronger pre-activation. Instead, it demonstrates the presence of a prior-driven effect, which intentionally triggers the modulation of MEPs congruent with the prior, subsequently impacting the decision even at the trial-by-trial level.

Overall, the outlined effects demonstrated consistently through classical approaches such as trial-average up to trial-by-trial analysis, how motor cortices modulate their activity to facilitate the response congruent with the provided prior. These findings expand on previous research, highlighting that the influence of priors extends beyond the perceptual system to include the motor system.

However, within the sample, there was wide inter-individual variability in the magnitude at which this effect manifests. Previous studies showed that the impact of prior knowledge strongly varies between individuals (Tarasi et al., 2022a; Tulver et al., 2019). Indeed, there exists a continuum of predictive strategies, ranging from individuals who shape external reality based on pre-established models to those who rely solely on current sensory data. Moreover, different predictive styles have been related to individuals’ dispositional factors such as autistic (ASD) and schizotypal traits (SSD) (Crespi and Badcock, 2008; Tarasi et al., 2022b). For instance, ASD individuals tend to rigidly rely on recent sensory information and bottom-up input (Pellicano and Burr, 2012; Tarasi et al., 2021; Ursino et al., 2022), while SSD traits emphasize prior experiences (Stuke et al., 2021; Tarasi et al., 2023b).

To date, no study has explored the presence of inter-individual difference in prior-driven motor system’s preparatory activity, nor whether ASD and SSD play a specific role in favoring these differences. First, we assessed whether the variability observed in prior-driven MEPs modulation was associated with different decision-making strategies. To this end, we employed a similar approach used by Tarasi et al., (2023b, 2022a), dividing the sample into two groups based on the degree of prior-dependent MEP modulation (ΔMEP_trial-averaged_). Analyses revealed that MEP_modulators_ (i.e., above-median) showed a significantly greater criterion modulation (Δcriterion = 1.22±0.21) compared to MEP_non-modulators_ (i.e., below-median; Δcriterion=0.46±0.13; t_60_=-3.09, p=0.003; BF=12.22), without any significant difference on sensitivity between the two groups (MEP_modulators_ vs MEP_non-modulators_: 0.99.±0.06 vs 1.08±0.07; p=0.350, BF=0.38). This finding corroborates the relationship between motor system activity and the weight assigned to prior expectations in decision-making. Therefore, MEP_modulators_ tend to adopt decisional strategies that are strongly influenced by prior expectation (*believers*), whereas MEP_non-modulators_ rely more on sensory information (*empiricists*), thus biasing less their decision according to expectations.

Finally, we questioned whether believers and empiricists correspond to cognitive styles spanning along the autism-schizotypy axis as proposed by Tarasi et al., (2022b). According to this model, believer’s and empiricist’s styles should be associated with schizotypal and autistic features, respectively. To identify where individuals lay on the autism-schizotypy axis, we conducted a principal component analysis (PCA) on the Autistic Quotient (Baron-Cohen et al., 2001) and Schizotypal Personality Questionnaire (Raine, 1991) subscales (Fig.S3). We extracted the first two principal components and selected the second one (PC2) for further analysis, as previous research found that PC2 effectively captures the diametric relationship between autistic and schizotypal traits(Dinsdale et al., 2013; Nenadić et al., 2021; Zhou et al., 2019). Subsequently, we assessed whether the two groups (*empiricists* and *believers*) exhibited different PC2 scores. Analyses revealed that *empiricists* exhibited negative PC2 values (−0.31±0.14), placing them closer to the autistic pole. Conversely, *believers* demonstrated positive PC2 scores (0.31±0.19), positioning them closer to the positive schizotypal pole of the continuum (t_60_=-2.57, p=0.013; BF=3.92). These findings corroborate predictive coding interpretation of ASD and SSD, demonstrating that SSD traits are associated with predictive strategy that overweight prior information, whereas ASD traits are linked to a more sensory-driven decision-making approach(Tarasi et al., 2022b). Mediation analysis corroborated this claim, demonstrating that the position along the SSD-ASD continuum modulates the decision bias adopted by the participants. Importantly, this effect was fully mediated by the concurrent modulation of M1s activity (Fig.4B). This suggests that participants positioned closer to the SSD vs. ASD pole of the continuum exhibited a strong vs. weak modulation of the decision criterion through a substantial vs. dampened shift of motor excitability following prior induction.

**Figure 4.**
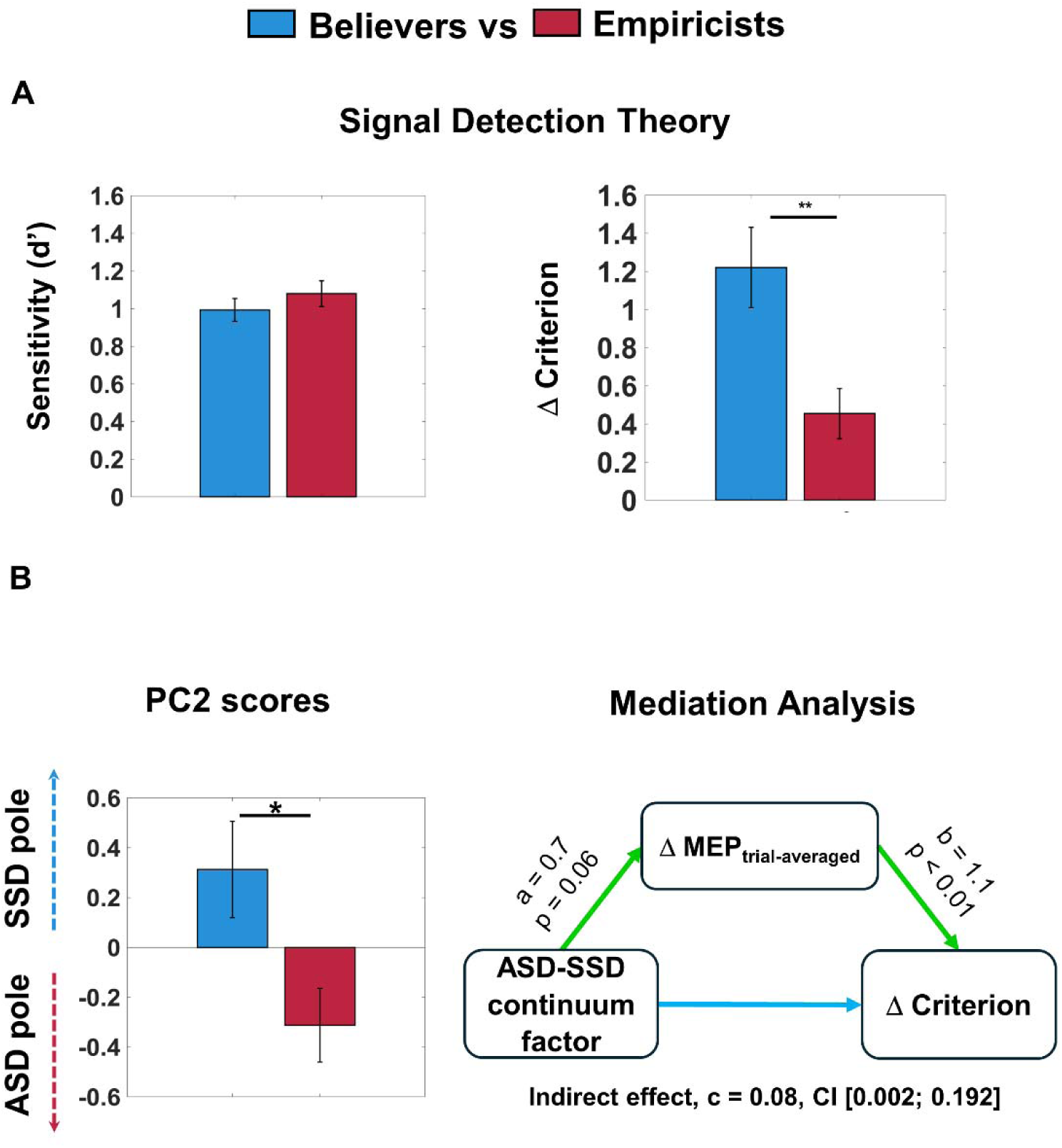
Empiricists and Believers adopt different decisional strategies and lay on opposite poles of the ASD-SSD continuum. **A.** SDT indices of Sensitivity (d’) and Criterion (c) are represented separately for Empiricists and Believers. The two groups were equal in sensitivity values, while prior-dependent shift in criterion was significantly greater in the believers vs. empiricists group. Data are represented as mean ± SEM; **p<0.01. **B.** PC2 scores represent a proxy of individuals’ position along the ASD-SSD continuum. Positive vs. negative values are associated with schizotypal vs. autistic traits, thus indicating proximity of believers and empiricists to the SSD and ASD pole, respectively. Mediation analysis corroborated this claim, demonstrating that the position along the SSD-ASD continuum modulation on the decision bias adopted by the participants, with this influence being fully mediated by the concurrent modulation of M1s activity. This suggests that participants positioned closer to the SSD vs. ASD pole of the continuum exhibited a strong vs. weak modulation of the decision criterion through a substantial vs. dampened shift of motor excitability following the prior induction. Data are represented as mean ± SEM; *p<0.05.

## Discussion

The presented study demonstrates the crucial role of the motor system in encoding decisional priors. Specifically, TMS delivered to the M1s areas elicited higher vs. lower MEPs when there was a congruency vs. incongruency with the prior-related action and trial-by-trial fluctuations in MEPs amplitude intercepted the prior-driven decisional bias.

These results shed lights on the instantiation of predictive strategies within the motor system, highlighting and extending a more general asset which we previously observed within the perceptual system, where perceptual priors were able to shape behavior through a modulation of alpha oscillations amplitude (Kloosterman et al., 2019; Tarasi et al., 2022a). Future investigation could explore the link between prior-driven mechanisms observed in the perceptual and motor system using combined TMS/EEG protocols to shed light on how the neural modulation driven by priors (alpha power) relates to the physiological modulation (MEPs) identified in the present study. One possibility is that these mechanisms align, suggesting a cohesive process between perceptual and motor domains. It is also plausible that they represent distinct mechanisms
subserving prior integration: there may be ‘motor individuals’ who just preprogram actions associated with priors, while ‘sensory individuals’ first prepare the perceptual areas to encode the actual stimuli by modulating alpha activity before engaging in motor processes.

An interesting result concerns the independence of the prior-effect on MEPs relative to the timing of TMS stimulation. One might have expected that, as participants approached the stimulus onset, the effect would have been more pronounced. However, priors were presented for 800ms, and TMS was administered only after its disappearance. This implies that, even with stimulation occurring furthest from the stimulus (i.e., −700ms), participants had already integrated the prior-related information and were preparing motor cortices activity accordingly. This result is particularly intriguing considering the nature of the paradigm used. While motor preparation is often studied in contexts where participants already have the information needed to make a choice (Duque et al., 2010; Greenhouse et al., 2015; Kennefick et al., 2019), in the employed scenario, participants had to take an additional step, the perceptual one, as it was necessary to analyze the moving dots to determine their direction. The strong effect identified demonstrates that even in these cases, participants begin to prepare their response before sensory analysis has occurred.

Moreover, our dual-coil paradigm allowed to investigate effects in the dominant vs. non-dominant hand (i.e., right vs. left), revealing distinct modulatory patterns. Overall, we observed a strong inhibition of the hand coding for the prior-incongruent action both in the right and in the left hand. However, we observed a heightened excitation of the congruent hand, but only when the non-dominant hand (i.e., left) coded for the prior-congruent response. This could imply the need to inhibit either hand when incongruent with expectations, while simultaneously exciting the non-dominant hand (i.e., left) given its comparatively lower motor proficiency. This suggests the need for introducing a supplementary mechanism for proper prior integration in this scenario. However, the involvement of both motor cortices in the inhibition process suggests its strong significance, potentially conferring an evolutionary advantage. Indeed, rather than overstimulating the cortex involved in the suggested movement, it reduces activity in the M1 cortex not coding for the prior-related action, thereby optimizing cognitive resources. These findings fit within theories conceptualizing predictive models as adaptive strategies essential for minimizing the “energetic cost” incurred by the cognitive system (Friston, 2009; Friston and Kiebel, 2009).

Furthermore, while the effects of priors on MEPs amplitude modulation were consistent, there was significant variability within our sample in the magnitude of this effect. This observation prompted us to investigate how these inter-individual differences might account for variations in decision-making strategy. Previous literature suggests that people vary along a continuum of predictive styles(Tarasi et al., 2023b, 2022a), with some individuals heavily relying on prior information (*believers*), while others prioritize sensory inputs over prior knowledge (*empiricists)*. We demonstrated that *empiricists* placed a weaker reliance on predictive mechanisms, maintaining motor cortex excitability in an unbiased state rather than pre-modulating it. In contrast, believers engaged strong bias mechanisms in the motor cortex, preparing for the prior-congruent response even before stimulus onset. This, in turn, results in a significant modulation of the decisional criterion.

Finally, we tested a recently proposed model according to which certain predictive styles may be closely related to personality traits along a continuum ranging from schizotypal (which favors a style based on priors) to autistic-like (which favors a style based on sensory evidence) traits (Tarasi et al., 2022b). Crucially, *empiricists* were characterized by a prevalence of autistic vs. schizotypal traits, whereas *believers* showed the reversed pattern. These findings align with the expanding literature focused on identifying the neuro-behavioral signatures associated with Bayesian processing and inter-individual differences in predictive mechanisms (Friedrich et al., 2022; Hein et al., 2023; Ivanova et al., 2024). Importantly, for the first time, we showed that individual’s position along the ASD-SSD continuum influences the motor strategies they adopt. This finding may provide mechanistical insight on certain characteristics observed along the spectra. For instance, SSD individuals often exhibit a “Jumping-to-conclusion” bias (Dudley et al., 2013; Falcone et al., 2015), prompting them to respond prematurely before accumulating sufficient decisional evidence. This mechanism could be underpinned by an over-facilitated motor preparation, leading to hastened action.

To sum up, the current work investigated the processes behind predictive perception showing how expectations are integrated into the human cognitive system. The findings reveal that prior knowledge shapes the decisional bias by modulating motor excitability, facilitating the implementation of prior-congruent responses. Furthermore, we highlighted individual differences in predictive styles, with *believers* and *empiricists* exhibiting hyper vs. lower modulation of motor cortex activity, along with specific personality traits across the ASD–SSD continuum. These results contribute to the broader understanding of how prior information guides perceptual processes, shedding light on the intricate interplay between expectations and motor preparation in shaping decision-making outcomes and in explaining the inter-individual differences in predictive processing.

## METHODS

### Participants

62 participants (37 female, age range 18-32) took part in the study. Before starting the experiment, they gave informed consent and were screened to avoid adverse reactions to TMS (Rossi et al., 2021). All the experimental procedures were performed in accordance with the Declaration of Helsinki.

### Autistic and Schizotypal traits

All participants were asked to complete two questionnaires: Autistic-Spectrum Quotient (AQ) (Baron-Cohen et al., 2001) and the Schizotipy Personality Questionnaire (SPQ) (Raine, 1991) to measure the autistic and schizotypal traits, respectively. Both AQ and SPQ are self-report questionnaires used to measure autistic and schizotypal traits in the non-clinical population. The AQ is composed of 50 questions divided in 5 subscales measuring five ASD features: imagination, communication, social skills, attention to detail, and attention switching. We applied the original scoring methods, converting each item into a dichotomous response (agree/disagree) and assigning a binary code (0/1) to the responses. The SPQ comprises 74 items divided in 9 subscales: ideas of reference, magical thinking, social anxiety, unusual perceptual experiences, constricted affect, no close friends, odd behavior, odd speech, and suspiciousness. We used the original scoring methods and assigned responses a binary code (0/1).

### Stimuli

Visual stimuli consisted of 400 white dots moving within a square region at the center of the screen (12.8° x 12°.8 visual angle) at a speed of 4.5°/second (Di Luzio et al., 2022). Stimuli were presented at a distance of 70 cm while participants sat comfortably in a dimly lit room using Matlab (version 2013b) and the Psychophysics toolbox. In each trial, dots could move with different levels of motion coherence (0%, 3%, 6%, 9%, 15%, 21%, 30%, or 60% of dots moving either to the left or to the right), determining different levels of task difficulty. 0% coherence means that all the dots move randomly; 80% coherence means that 320 dots move coherently rightward or leftward, while 80 dots move randomly. For the main task, motion coherence was set individually for each participant to the level at which they reached 70% of accuracy in the titration phase. The titration procedure is necessary to balance the difficulty level for each individual participant, preventing emerged results from being attributed to upstream factors linked to the ability to analyze incoming stimuli(Tarasi and Romei, 2023).

### Experimental protocol

Participants performed a random dot motion (RDM) discrimination task indicating the perceived direction of movement (i.e., rightward, or leftward) by pressing a key on the keyboard as quickly and accurately as possible. Responses required an abduction movement of the index finger to press a key on a reversed keyboard (F12 with the right hand for signaling rightward movement and F5 with the left hand for signaling leftward movement; see Fig.1A). EMG activity from the FDI muscle of both hands (agonist in index finger abduction) was recorded during the entire duration of the task.

In a preliminary phase, participants performed a brief and simplified demo version of the task with stimuli presented for 1000ms, and then a training phase in order to familiarize with the task. Then, participants underwent a titration procedure to determine the coherence of movement for which accuracy was 70%. Motion coherence thresholds were estimated for each participant using the method of constant stimuli and fitting the psychometric function with the *psignifit* toolbox (Fründ et al., 2011). Threshold values were defined as the coherence for which 70% accuracy was obtained.

The experimental phase comprised 4 test blocks of 120 trials each, where expectations about the upcoming stimulus were manipulated by presenting a probabilistic cue prior to stimulus presentation. In each test block, single-pulse TMS was delivered on both the right and left M1 at 4 different timings (−700 ms, −400 ms, −100 ms, +200 ms relative to stimulus onset) at 110% of the previously determined rMT. Pulses were delivered with an Inter-pulse Interval (IPI) of 1ms, to avoid physical interactions between the coils (Cincotta et al., 2005; Grandjean et al., 2018). Each trial started with a fixation point (3.5 s), then a probabilistic cue was presented for 1 s (left/right arrow or cross). After a delay, the random-dot stimulus was presented at the previously determined motion coherence threshold at the center of the screen for 400ms. Participants had to report the direction of movement by pressing the corresponding key on the keyboard as accurate and fast as possible. Trials with reaction times (RT) below 200ms were excluded from data analysis to avoid considering trials where participants responded before the TMS pulse was delivered in the +200 condition. Cues were either informative (right or left arrow) or uninformative (cross) and indicated the probability of rightward or leftward movement. The arrow pointing to the right (vs. left) indicated a high probability (70%) of rightward (vs. leftward) movement, while the cross indicated an equal probability (50%) for either movement directions. Participants were explicitly informed that the probabilistic cue was valid (i.e., congruent with the actual probability).

### Behavioural data analysis

#### Signal Detection Theory (SDT)

We computed SDT indices d’ and c(Green and Swets, 1966) to evaluate participants’ sensitivity (higher d’ values indicate higher sensitivity) and criterion (c different from 0 indicates the presence of response bias). These indices were calculated based on the proportion of hits (i.e., responding right in rightward trials) and false alarm (i.e., responding left in rightward trials). To investigate the effect of prior information on sensitivity and criterion, we computed d’ and c separately for trials preceded by left, right or neutral cues. We conducted an ANOVA on both c and d’ with cue type as the within factor (3 levels: left, right, neutral).

#### Neurophysiological assessment

The experiment started with electrode montage setup, detection of optimal scalp position and assessment of resting motor thresholds (rMT). AG/AgCl surface electrodes were positioned in a belly-tendon montage over the right and left first dorsal interosseus muscle (FDI). Electromyographic (EMG) signals were recorded using a Biopac MP-35 (Biopac, U.S.A) electromiograph, band-pass filtered between 30 and 500 Hz and sampled at 10 kHz. TMS was performed using a Magstim Bistim² stimulator. Pulses were delivered through a 50mm figure-of-eight coil and were remotely triggered by a MATLAB script (The MathWorks, Natick, USA). Left and right M1 were localized as the optimal scalp position where MEPs of maximal amplitudes could be induced in the contralateral FDI. Coils were positioned over M1, tangentially to the scalp and with a 45° angle from the midline, to induce a posterior-anterior current flow in the brain(Di Lazzaro et al., 2004). We first determined the individual rMT, defined as the minimum stimulator output intensity that evokes MEPs with an amplitude of at least 50 mv in 5 out of 10 consecutive trials (Rossini et al., 1994). Then, we assessed the intensity required to elicit MEPs with an average peak-to-peak amplitude of 1 mV.

#### Neurophysiological data processing

Neurophysiological data were processed offline. MEP peak-to-peak amplitudes were measured in the 15-60ms window following the TMS pulse. We discarded MEPs showing EMG activity above 0.3 mV in the preceding 350ms time-window and removed trials with MEPs below 0.1 mV (Dupont-Hadwen et al., 2019; Klein-Flügge et al., 2013), leading to a 12% of trials removed. As a control analysis, we replicated this procedure with more liberal criteria, discarding only trials showing precontractions above 0.3 mV (1% of trials removed). This additional analysis revealed the same pattern of results observed with more conservative MEPs removal criteria (see Supplementary Materials). To investigate the influence of prior information on EMG activity, and whether this effect was modulated by TMS timing and differed between the two hands, we conducted a three-way ANOVA on MEPs amplitude. We considered prior information (3 levels: right, neutral, left), time (4 levels: −700, −400, −100, +200 ms) and hand (2 levels: right, left) as within factors. To further explore the results of the ANOVA, post-hoc analyses were conducted using paired-sample t-tests.

#### Brain-to-behavioral analysis

To investigate whether there was an association between behavioral and neurophysiological data, we computed two indices representing the prior-dependent modulation of criterion and the prior-dependent modulation of MEP activity. For behavioral data, we took the difference between criterion values in the right vs left prior conditions (Δ criterion = c_right – c_left). This index represents how much individuals shift their criterion between cue-right and cue-left conditions, thus measuring the influence of the cue on participants’ choices. For EMG data, we first computed ΔMEP separately for the two hands by subtracting MEP activity in the prior congruent vs incongruent conditions (ΔMEP right hand = MEP amplitude with the right arrow – MEP amplitude with the left arrow; ΔMEP left hand = MEP amplitude with the left arrow – MEP amplitude with the right arrow). Then, we summed these two values to obtain a global measure of prior-dependent MEPs modulation (ΔMEP_trial-averaged_= ΔMEP_right_ + ΔMEP_left_). We then conducted Pearson and Spearman correlation analyses considering Δ criterion and ΔMEP_trial-averaged_. To verify the robustness of the association, we run the skipped correlation (both Pearson and Spearman) using the Robust Correlation toolbox (Pernet et al., 2013).

#### Trial-by trial MEPs – starting point

We used a Drift Diffusion Model (DDM) to investigate the trial-by-trial relationship between MEPs and the decisional bias of the participant. The drift diffusion model (DDM) is a computational model commonly used in cognitive psychology and neuroscience to understand decision-making processes. It posits that decision-making involves accumulating evidence over time until a decision threshold is reached (Ratcliff et al., 2016). This model allows to extract several decision-making parameters, each intercepting the different component leading to a particular choice. The drift rate parameter (v) represents the rate at which evidence accumulates in favor of one decision option over the other. A higher drift rate indicates faster and more accurate decision-making. The threshold parameter (a) represents the amount of evidence needed to make a decision. Higher thresholds lead to more cautious decision-making, while lower thresholds result in quicker decisions. The bias parameters (z) reflect the starting point of evidence accumulation towards one decision option over the other. A bias towards one option (e.g., rightward motion) means that evidence starts accumulating in favor of that option from the beginning of stimulus presentation. Based on the previous results derived from the SDT, showing a strong effect of both priors and MEPs on decisional bias, we set up two regression-based drift diffusion model (Wiecki et al., 2013) to assess whether the trial-by-trial MEPs fluctuations predicted the starting point of the accumulation process.

The initial model aimed to determine whether the trial-by-trial variability between congruent and incongruent MEPs influenced participants towards congruent or incongruent decisions. First, we recoded both the presented stimulus and the participant’s response as either congruent or incongruent. A congruent stimulus/response occurred when the current stimulus/response aligned with the prior expectation (e.g., following a prior indicating rightward movement: a right stimulus was congruent; a left stimulus was incongruent; the same applied to the response). Following this, we recoded z-scored MEPs in a similar fashion: MEPs were considered congruent if they originated from the hand corresponding to the prior-congruent response, and vice versa for incongruent MEPs. Then, we calculated the difference between congruent and incongruent MEP (delta MEP_trial-by-trial_) to derive a single index suitable for fitting the DDM. Finally, we applied a drift diffusion model using this recoding of stimulus and response. We introduced the delta MEP_trial-by-trial_ as a regressor to assess its potential impact on the starting point parameter. We hypothesized that an increase in MEP in the congruent hand compared to the incongruent hand should bias participants towards a congruent response that aligns with the provided prior information, and vice versa.

The second model, on the other hand, did not account for the congruence factor. Instead, it utilized the stimulus and the actual response presented to the participant in each trial as inputs. As a regressor, this time, the difference between z-scored MEPs extracted from the right hand versus the left hand was employed. Therefore, a positive value indicated that, in that specific trial, the MEP recorded from the right hand was greater than the MEP recorded from the left hand. Additionally, we evaluated whether the slope differed depending on the 3 different priors presented. The logic of the analysis is to assess whether a potential larger amplitude of the MEP in the right hand (vs. the left hand) would push the participant to shift the starting point closer to the decision threshold corresponding to the rightward (vs. leftward) choice. Being a hierarchical Bayesian models, we assessed the significance of the regressor extracted in the first model (delta MEP_trial-by-trial_) and the three regressors extracted in the second model (i.e., one for each cue) by evaluating whether the posterior distribution overlapped with zero. If 95% of the distribution of the regressor does not overlap with zero, it implies that the predictor is significantly able to modulate the starting point parameter.

### MEP-modulators (Believers) vs MEP-unmodulators (Empiricists)

To explore whether interindividual differences in the prior-dependent modulation of MEP activity underpin different decision-making strategies, we divided our sample into two groups using a median split approach on ΔMEP, to mimic the approach used in Tarasi et al., (2023b, 2022a) separating individuals who showed large (vs low) MEP modulation in order to allow a direct comparison between studies. Specifically, we calculated the median of ΔMEP (median = 0.11) and extracted the following groups: MEP-modulators (i.e., individuals who show above-median ΔMEP) and MEP-unmodulators (i.e., individuals who show below-median ΔMEP). To investigate whether the two groups differed in their decision-making performance, we employed an independent samples t-test to evaluate differences in either sensitivity or Δcriterion between groups.

### PCA and Mediation Analysis

Based on previous evidence indicating the impact of personality traits, specifically the presence of autistic vs schizotypal traits, on decision-making strategies (Andersen, 2022; Brosnan et al., 2014; Tarasi et al., 2023b, 2023a), we analyzed participants’ responses to the Autism-Spectrum Quotient (AQ) and Schizotypal Personality Questionnaires (SPQ) and their relationship with decision-making strategies. First, we identified individuals’ position on the autism-schizophrenia axis through Principal Component Analysis (PCA), which was conducted on the correlation matrix of the AQ and SPQ subscales. First, we confirmed the feasibility of the PCA using the Kaiser-Mayer-Olkin (KMO) and the Bartlett’s test. We then extracted the first two components and conducted further analysis on the second one (PC2), which is expected to exhibit opposite loadings with autism and schizotypy, as reported by several previous studies (Dinsdale et al., 2013; Nenadić et al., 2021; Tarasi et al., 2023b; Zhou et al., 2019). By analyzing the correlation between Principal Component and subscales we further confirm the presence of this diametrical pattern (See supplementary materials). Subsequently, individual scores along PC2 were extracted, representing a proxy of participants’ position along the ASD-SSD continuum. Finally, to assess whether individual’s position along this axis could explain interindividual differences in the predictive style adopted, we conducted an independent-sample t-test on individual PC2 score to determine if MEP-modulators and MEP-unmodulators showed significantly different scores. Moreover, to further investigate the crucial role of ASD-SSD traits in dictating decision-making strategies we have conducted a mediation analysis (PROCESS, SPSS). PC2 scores was used as a prediction of the delta criterion, evaluating whether the delta MEP mediated this relationship. The logic underlying this analysis is that we expect that the position along the continuum could shape decisional criterion by means of a concurrent modulation of the physiological marker.

## Notes

### Competing Interest Statement

The authors have declared no competing interest.

