## Supplementary Materials for "Preparing to act follows Bayesian inference rules"

**Time effect on MEPs amplitude**

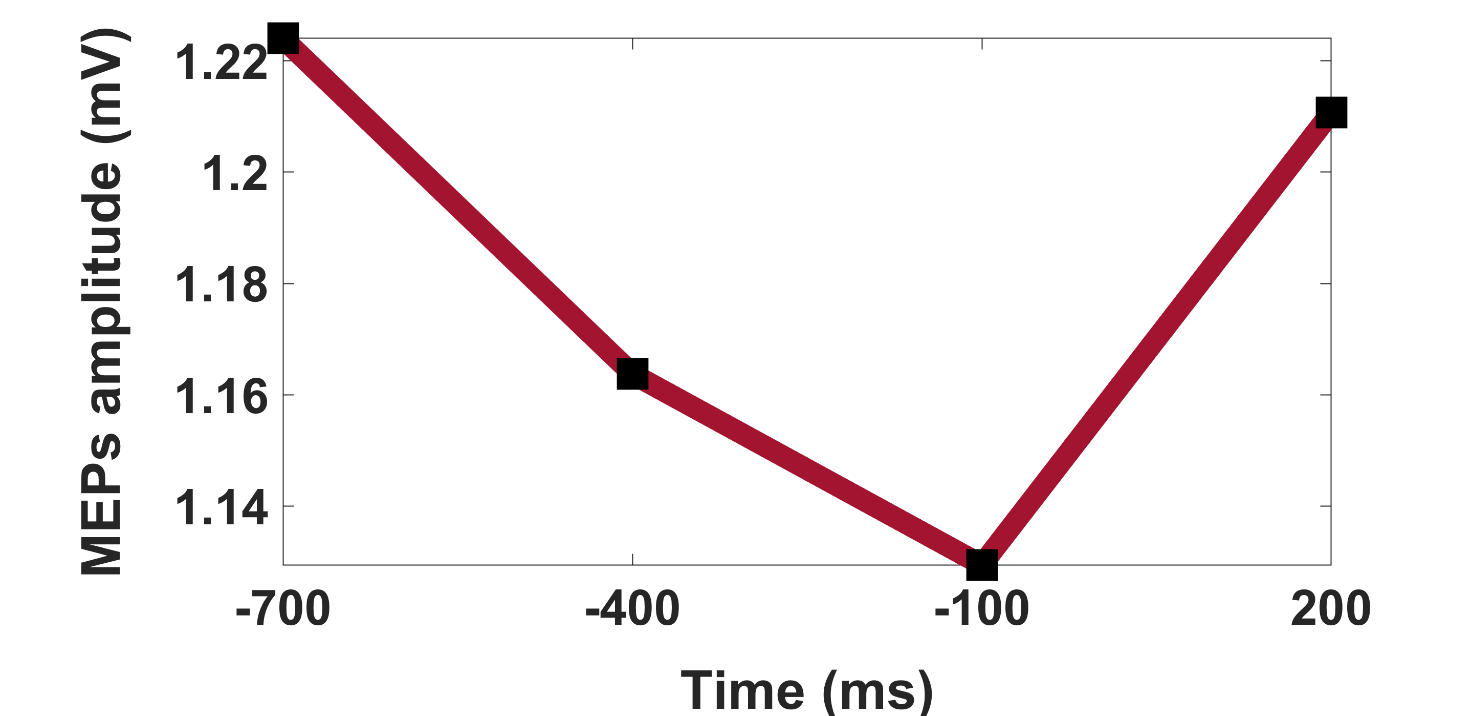

**Figure S1**

*As discussed in the main text, we conducted a three-way ANOVA on MEPs amplitude to investigate the influence exerted by prior expectations on motor system excitability. Specifically, we considered prior information (3 levels: right, neutral, left), time (4 levels: -700, -400, -100, +200 ms) and hand (2 levels: right, left) as within factors. We found a significant interaction between hand and prior, indicating different prior-dependent modulations between the two hands. Furthermore, there was a significant effect of TMS timing on MEPs activity (F_3,183_=14.23, p<0.001), but any significant interaction with the other factors considered (i.e., hand and cue), meaning that MEP amplitude changed as a function of time, but the effect was the same regardless of the presented cue or the considered hand. For this reason, we decided to collapse the time factor in subsequent analyses and considered the four TMS conditions (-700ms, -400ms, -100ms, +200ms) altogether. Here we have represented the main factor of time, which demonstrates a progressive decrease in cortical excitability as the stimulus approaches, followed by a rise when the stimulus is presented.*

**Adopting More Liberal Criteria for MEPs Removal Does Not Affect Results**

Processing of electromyographic (EMG) included the removal of MEPs showing precontractions above 0.3 mV in the preceding 350ms and MEPs below 0.1 mV, which led us to a 12% of trials removed. This widely employed strategy allows for the elimination of trials where it is unclear whether an MEP has been elicited or if the recorded activity is merely due to stochastic fluctuations in electromyographic activity

As a control analysis, we applied more liberal criteria, discarding only MEPs showing precontractions above 0.3 mV in the preceding 350ms before TMS pulse. This procedure led to a 1% of trials removed. We replicated the three-way ANOVA on MEPs amplitude considering the remaining trials. As reported in the STAR Methods section, we considered prior information (3 levels: right, neutral, left), time (4 levels: -700, -400, -100, +200 ms) and hand (2 levels: right, left) as within factors. This control analysis revealed the same pattern of results, showing a significant interaction between hand and prior (F2,122=13.79; p<0.001) and a main effect of time (F2,122=17.12; p<0.001). Post-hoc analyses indicated the presence of a congruency effect between hand and prior. In the right hand, MEPs were higher vs. lower when the rightward (1.20±0.07) vs. leftward (1.08±0.06, t61=--2.92; p=0.005, BF=6.39) cue was presented. With the neutral prior, MEPs were higher (1.19±0.07) relative to the leftward condition (t61=-3.34; p=0.001; BF=19.23) but not significantly different from the rightward condition (t61=-0.53; p=0.60; BF=0.16).

As for the left hand, higher activity was found when the leftward prior was presented (0.98±0.06) relative to when the rightward prior was presented (0.90±0.06; t61=4.20; p<0.001; BF=238.57). In the neutral condition (0.94±0.06), MEPs were higher than in the rightward condition (t61=3.36; p<0.001; BF=20.14) and lower than in the leftward condition (t61=2.30; p=0.025; BF=1.60). The reported analyses highlight the same pattern of results observed when adopting more conservative criteria for MEPs removal, thus validating the results from our first analyses. Crucially, the same pattern of results emerges when we consider only the three pre-stimulus timings (i.e., -700, -400, -200) in the ANOVA, whether using the conservative or liberal criteria for MEP removal.

**Removing the data from the TMS pulse delivered after the stimulus presentation does not affect the results.**

Our paradigm consisted of four stimulation conditions, where TMS pulses were delivered at different timings relative to stimulus onset (-700, -400, -100, +200ms). As a control analysis, we investigated MEPs amplitude modulations considering only pre-stimulus conditions, thus removing from data analysis trials in which the TMS pulse was delivered 200ms after stimulus onset. We replicated the three-way ANOVA on MEPs amplitude with prior information (3 levels: right, neutral, left), time (3 levels: -700, -400, -100ms) and hand (2 levels: right, left) as within factors. The ANOVA showed a significant interaction between hand and cue (F_2,122_=12.39; p<0.001) and a main effect of time (F_2,122_=17.36; p<0.001; See Fig.S1 for graphic representation of the temporal evolution in MEPs amplitude). We further explored the interaction hand*cue using paired sample one-tailed t-tests. The right hand showed higher vs. lower MEPs when the rightward (1.30±0.07) vs. leftward cue (1.20±0.06; t_61_=2.78; p=0.004) was presented. With the neutral cue, MEPs were higher (1.31±0.07) relative to the leftward condition (t_61_=3.27; p<0.001) but not significantly different from the rightward condition (t_61_=-0.35; p=0.64). The left hand showed higher vs. lower MEPs when a leftward (1.12±0.06) vs. rightward cue (1.02±0.05; t_61_=4.16; p<0.001) was presented. The neutral cue led to higher MEPs (1.09±0.06) relative to the rightward cue (t_61_=3.62; p<0.001) and lower MEPs relative to the leftward cue (t_61_=1.88; p=0.033). These results corroborate our initial analysis, indicating that the hand-dependent effect of priors on MEP activity described in the main text was not guided by a post-stimulus process.

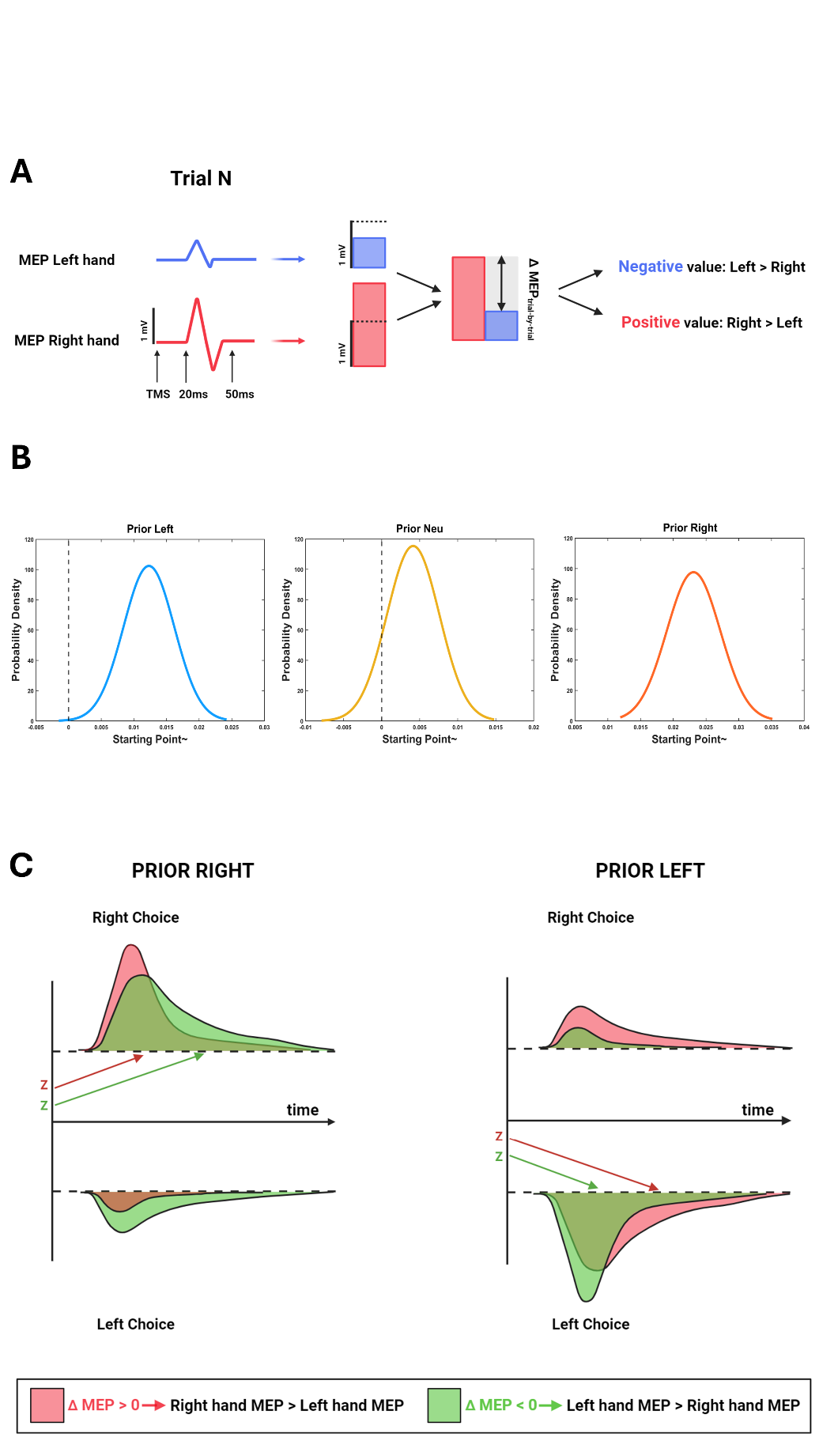

**Figure S2**

***A****. The second DDM model confirmed the results of the first one (see Main text). Indeed, it demonstrated that the model 1 results remained valid even after removing the congruency effect. Specifically, it showed that trials where the MEPs of the right hand were relatively larger in amplitude than those of the left hand were associated with a stronger rightward decisional bias. Conversely, when the MEPs of the left hand exceeded those of the right hand, there was a bias towards reporting leftward movement.*

***B****. Crucially, this effect interacted with the provided priors. In conditions where the prior was informative (i.e., right and left), the significant effect persisted (p<0.001). However, in conditions where the prior was uninformative, MEP fluctuations did not predict the shift in the starting point parameter (p=0.12).*

***C****. This suggests that the trial-by-trial effect is not merely a stochastic mechanism, where the decision shifts based on which M1 area exhibits stronger pre-activation. Instead, it demonstrates a prior-driven effect, which intentionally triggers the modulation of MEPs congruent with the prior, subsequently impacting the decision even at the trial-by-trial level.*

|  | **Component Loadings** | |
| --- | --- | --- |
|  | **PC1** | **PC2** |
| AQ – Social skills | 0.712 | **-0.360** |
| AQ – Attention switching | 0.525 | **-0.438** |
| AQ – Attention to detail | 0.257 | 0.655 |
| AQ – Imagination | 0.192 | **-0.391** |
| AQ – Communication | 0.664 | -0.090 |
| SPQ – Reference | 0.449 | **0.369** |
| SPQ – Magical thinking | 0.311 | **0.779** |
| SPQ – Social anxiety | 0.644 | -0.343 |
| SPQ – Unusual perceptual experiences | 0.581 | **0.592** |
| SPQ – Odd behavior | 0.672 | 0.312 |
| SPQ – No close friends | 0.759 | -0.156 |
| SPQ – Odd speech | 0.568 | 0.253 |
| SPQ – Constricted affect | 0.595 | -0.271 |
| SPQ – Suspiciousness | 0.603 | -0.252 |

**Figure S3**

*Principal component analysis isolated two components that together explain 53% of the variance. The questionnaire loadings are consistent with those identified in the literature, with the first component showing a positive correlation with all subscales, collapsing the common information between the two conditions. The second component, on the other hand, traces the ASD-SSD continuum, having positive loadings with the positive subscales of the SPQ and negative loadings with the subscales of the AQ.*
